# *RAB18* is a key regulator of GalNAc conjugated siRNA induced silencing in Hep3B cells

**DOI:** 10.1101/2022.01.07.475411

**Authors:** Jiamiao Lu, Elissa Swearingen, Miki Hardy, Patrick Collins, Bin Wu, Eric Yuan, Daniel Lu, Chi-Ming Li, Songli Wang, Michael Ollmann

**Affiliations:** Genome Analysis Unit, Amgen Global Research, South San Francisco, USA, 1120 Veteran Blvd, ASF1, South San Francisco, CA94080

**Author notes:** Correspondence Authors: Jiamiao Lu PhD, Michael Ollmann PhD.

**Keywords:** RAB18, GalNAc conjugated siRNA, siRNA silencing efficacy, siRNA delivery, pooled genome wide knockout screen

## Abstract

Small interfering RNAs (siRNA) therapeutics have developed rapidly in recent years, despite the challenges associated with delivery of large, highly charged nucleic acids. Delivery of siRNA therapeutics to the liver has been established, with conjugation of siRNA to N-acetylgalactosamine (GalNAc) providing durable gene knockdown in hepatocytes following subcutaneous injection. GalNAc binds the asialoglycoprotein receptor (ASGPR) that is highly expressed on hepatocytes and exploits this scavenger receptor to deliver siRNA across the plasma membrane by endocytosis. However, siRNA needs to access the RNA-induced silencing complex (RISC) in the cytoplasm to provide effective gene knockdown and the entire siRNA delivery process is very inefficient, likely due to steps required for endosomal escape, intracellular trafficking, and stability of siRNA. To reveal the cellular factors limiting delivery of siRNA therapeutics, we performed a pooled, genome wide knockout screen based on delivery of GalNAc conjugated siRNA targeting the *HPRT1* gene in the human hepatocellular carcinoma line Hep3B. Our primary pooled genome wide knockout screen identified candidate genes that when knocked out significantly enhanced siRNA efficacy in Hep3B cells. Follow-up studies indicate that knockout of one gene in particular, *RAB18*, improved siRNA efficacy.

## INTRODUCTION

SiRNAs are short (20~25 base pairs), double-stranded RNA molecules that operate through the RNA interference (RNAi) pathway to specifically degrade target gene mRNA^1^. The therapeutic potential of siRNAs has been intensively investigated in recent years to treat a wide range of human diseases. Compared with traditional drug molecules, siRNAs are highly potent and capable to act on previously “non-druggable” targets^2–4^. More impressively, the duration of siRNA conjugate mediated potent mRNA knockdown has been shown to last for several months^2–5^.

Despite their substantial therapeutic potential, siRNA therapeutics are associated with challenges associated with the delivery of large, highly negatively charged nucleic acids into cells. Delivery of siRNA therapeutics to the liver has been established, with conjugation of siRNA to GalNAc providing durable gene knockdown in hepatocytes following subcutaneous injection^6–8^. On hepatocytes, GalNAc binds the highly expressed scavenger receptor, ASGPR, to deliver siRNA across the plasma membrane by clathrin-coated endosomes^9–11^. The human ASGPR exists as hetero-oligomers formed by two subunits: the major ASGR1 (asialoglycoprotein receptor 1) subunit and the minor ASGR2 (asialoglycoprotein receptor 2), with the ASGR1 subunit being critical for efficient GalNAc conjugated siRNA delivery^12–14^. Although GalNAc conjugation improves siRNA delivery, it is still an inefficient process^15,16^.

As endosomes mature, the internal pH drops and causes GalNAc conjugated siRNAs to be released from ASGPR. The ASGPR receptors then quickly recycle back to the cell surface, while GalNAc conjugated siRNAs remain inside the endosome^15^. The endosomal glycosidase then works to cleave GalNAc from siRNA conjugates^15^. Less than 1% of the remaining free siRNAs are capable of escape from endosomes through an unknown mechanism and have access to RISC in the cytoplasm to provide effective gene knockdown and induce RNAi responses in the cytoplasm^16^. After siRNA enters the cell, it remains inactive until becomes loaded into the core component of RISC. The passenger (sense) strand is cleaved and ejected at Argonaute 2 (Ago2), and the guide (antisense) strand is then bound to catalytic Ago2^17,18^. The siRNA guide strand then guides and aligns the RISC complex on the target mRNA and induces cleavage of the target mRNA through the catalytic function of Ago2. The siRNA intracellular trafficking and escape steps are very inefficient, and the underlying mechanisms are not fully understood^15,19^.

In recent years, adaptation of the bacterial CRISPR-Cas9 (Clustered Regularly Interspaced Short Palindromic Repeats-CRISPR-Associated Protein 9) system to mammalian cells have enabled genome wide loss-of-function screens to identify new biological mechanisms^20–24^. To reveal the cellular factors limiting delivery of siRNA therapeutics, we performed a pooled, genome wide CRISPR-Cas9 screen (referred as CRISPR screen in the rest of this article) based on delivery of GalNAc conjugated siRNA targeting the *HPRT1* gene in the human hepatocellular carcinoma line Hep3B. Multiple candidate genes that when knocked out significantly enhance siRNA efficacy in Hep3B cells were identified from the CRISPR screen. A secondary, arrayed CRISPR screen using multiplexed synthetic gRNA in 96/384-well format was then used to validate these candidate genes. Additional follow-up studies of one top candidate gene, *RAB18*, indicate that knocking out *RAB18* improves siRNA silencing potency at the mRNA level. The results of this study provide insights into mechanisms of siRNA delivery to both hepatic and extrahepatic tissues.

## RESULTS

### Hep3B cells demonstrated robust GalNAc conjugated siRNA induced silencing

An ideal system for identifying key regulators of GalNAc conjugated siRNA induced silencing would have the following attributes: 1) long-term maintenance; 2) stable Cas9 expression; 3) capability of gRNA lentivirus library transduction; and 4) sufficient siRNA induced silencing to allow ranking of candidate genes. Human primary hepatocytes have been proven to uptake GalNAc conjugated siRNA through cell surface ASGPR^9–11^. However, large scale CRISPR screens have been challenging in human primary hepatocytes due to their limited proliferative potential. We therefore explored the possibility of using human hepatocellular carcinoma cell lines such as HepG2 or Hep3B to perform our CRISPR screen. Although both HepG2 and Hep3B cells express high levels of *ASGR1* and *ASGR2* (Supplementary Table 1), only Hep3B cells displayed robust knockdown of target genes through GalNAc conjugated siRNA induced silencing in our hands (Figure 1a, Supplementary Table 2). GalNAc conjugated siRNA was able to induce target gene knockdown in Hep3B cell line in a dose dependent manner, and the level of silencing was sufficient to perform the CRISPR screen.

**Figure 1.**
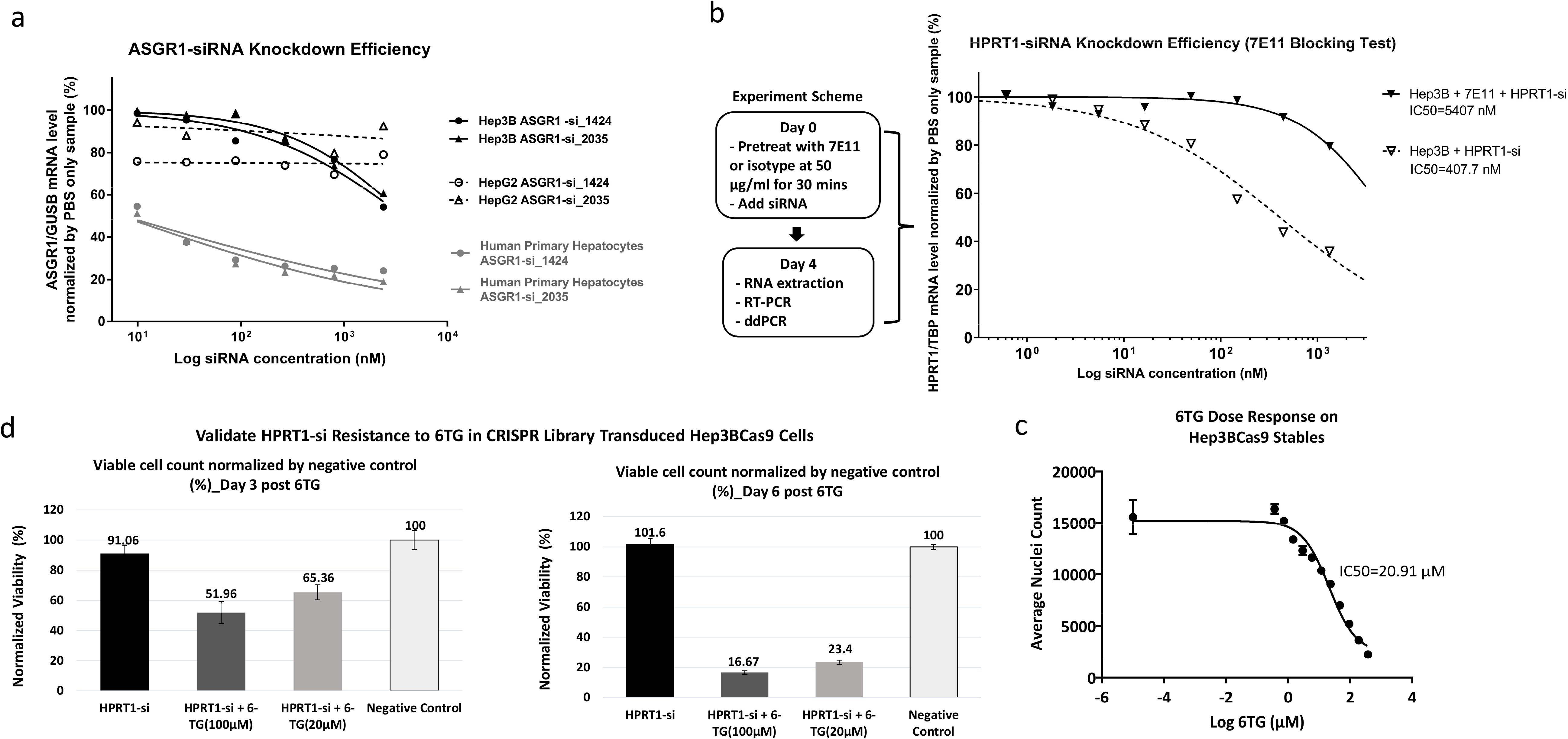
Validation of screen conditions for pooled genome wide CRISPR-Cas9 screen. a) Comparison of target gene (*ASGR1*) silencing potency in human primary hepatocytes, Hep3B and HepG2 cells by two GalNAc conjugated *ASGR1* siRNAs. The features of these two siRNA conjugates are described in Supplementary Table 2. b) Treatment with an in-house made anti-ASGR1 antibody, 7E11, mitigated the *HPRT1* gene silencing induced by GalNAc-*HPRT1* siRNA (8172) in Hep3B cells. Left panel outlines the experiment scheme and the right panel shows the ddPCR measurement of *HPRT1* mRNA levels in percentage normalized by housekeeping gene *TBP* readings and no siRNA (PBS only) treated control group. The feature and sequence of siRNA 8172 is described in Supplementary Table 2. c) Dosage dependent kill curve of 6TG treatment in Hep3BCas9 cells. d) A small-scale pilot experiment to validate the feasibility of using HPRT1-6TG live/dead selection for CRISPR screen. The gRNA lentivirus library transduced Hep3BCas9 cells were treated with GalNAc-*HPRT1* siRNA and/or 6TG (100 μL) in different groups. The viable cell count measured by ViCell on day 3 and day 6 post-6TG treatment for each treatment group was normalized by negative control group readings. The resulting normalized viability percentage of each group at both time points was plotted into bar graph. Left panel: Day 3 post-6TG treatment data. Right panel: Day 6 post-6TG treatment data.

To further validate whether GalNAc conjugated siRNA induced silencing in Hep3B is mediated through ASGR1, an antibody blocking test was performed (Figure 1b). To perform this experiment, Hep3B cells were first pre-incubated with an in-house generated, anti-ASGR1 antibody (7E11), or no antibody treatment as control for half an hour, followed by treatment with GalNAc conjugated siRNA targeting *HPRT1* (GalNAc-*HPRT1* siRNA: 8172) (Supplementary Table 2) at multiple doses. The target gene (*HPRT1*) mRNA levels were measured on day 4 post-siRNA treatment through ddPCR (digital droplet polymerase chain reaction) analysis. As indicated in Figure 1b, application of the ASGR1 specific antibody was able to mitigate the siRNA silencing efficacy (13-fold higher IC50 for no antibody (5404nM) relative to anti-ASGR1 (407.7nM).

After establishing the suitability of Hep3B for GalNAc conjugated siRNA induced silencing, we then generated Hep3B cells stably expressing Cas9. The editing capability of the Cas9 stable Hep3B was assayed through validating their editing efficacy on two target genes, *SLC3A2* and *ASGR1* (Supplementary Table 3 and Supplementary Figure 1). The Cas9 stable Hep3B cells (referred as Hep3BCas9 in the rest of this article) were then used to perform the CRISPR screen to search for key regulators of GalNAc conjugated siRNA induced silencing.

### HPRT1-6TG (6-ghioguanine) live/dead selection based CRISPR screen in Hep3BCas9 cells

A CRISPR knockout screen based on live/dead selection is the most efficient and convenient systematic experiment to identify potential regulators of siRNA efficacy. We therefore chose to take advantage of the established HPRT1-6TG based live/dead selection system for this CRISPR screen to look for key regulators of GalNAc conjugated siRNA induced silencing. 6-thioguanine (6TG), a purine analog, is incorporated into DNA and RNA resulting in cell death after being phosphorylated by hypoxanthine phosphoribosyl transferase (HPRT) encoded in humans by the *HPRT1* gene^25^. Knocking down or knocking out *HPRT1* provides resistance to 6TG and allows those cells to survive. One thing to keep in mind when utilizing HPRT1-6TG live/dead selection system is that mismatch repair defective cells and cells with *HPRT1* mutations might also be resistant to 6TG^26^, necessitating independent validation of candidates identified using this method. Based on this theory, a GalNAc conjugated siRNA targeting human *HPRT1* incorporating Fluoro (F) and Methoxy (OMe) modifications (Supplementary Table 2), GalNAc-*HPRT1* siRNA (8172), was designed and validated. If GalNAc-*HPRT1* siRNA can enter the cells and induce *HPRT1* gene silencing, these cells would be able to survive in the presence of 6TG. Otherwise the cells would be killed by 6TG selection if GalNAc-*HPRT1* siRNA does not silence *HPRT1*. Under the CRISPR knockout condition, if a gene is normally required for siRNA activity, knocking out this gene would diminish or abolish siRNA function and cause the cells to be eliminated by 6TG selection. Alternatively, if a gene normally functions to inhibit or block siRNA activity, knocking out this gene would improve siRNA potency and enable the cells to survive 6TG selection. Therefore, when sequencing gRNAs in surviving cells, the enriched gRNAs reflect genes that may normally inhibit siRNA activity, while gRNAs targeting genes essential for siRNA function would be depleted. However, other gRNAs targeting genes that impact cell viability through non-siRNA related mechanisms would also be depleted from live cell population, making it difficult to identify siRNA essential genes from the depleted gRNA population. We therefore chose to focus on analyzing the enriched gRNAs from surviving cells to enable us to identify genes that inhibit GalNAc conjugated siRNA induced silencing in Hep3B.

We first established the baseline 6TG kill curve in Hep3BCas9 cells without siRNA treatment (Figure 1c). To avoid both insufficient and excessive killing caused by 6TG, we performed a small-scale pilot run using 100 μM 6TG (~IC70) (Figure 1d) and 20 μM 6TG (~IC50) (Supplementary Figure 2). An 80k genome wide CRISPR gRNA lentivirus library (CRISPR KOHGW 80K (lot#17050301), Cellecta, Mountain View, CA) was transduced into Hep3BCas9 cells to generate a genome wide knockout pool. These gRNA transduced cells were then analyzed for their ability to be selected using GalNAc-*HPRT1* siRNA and 6TG. As illustrated in Figure 1d, cells were divided into four groups (0.6E+06 cells/group): 1) siRNA only, 2) siRNA with 6TG treatment, 3) 6TG only, and 4) negative control. To obtain sufficient but not excessive siRNA effect, 750 nM (about IC60) GalNAc-*HPRT1* siRNA (8172) was used. On day 3 post-6TG treatment, the 6TG only group had 35% viable cells while the *HPRT1*-si + 6TG group had 52% viable cells (Figure 1d). On day 6 post-6TG treatment, the 6TG only group had only 5% viable cells while the *HPRT1*-si + 6TG group had 17% viable cells (Figure 1d). These results indicate that GalNAc-*HPRT1* siRNA treatment was partially protective. This provides a screening phenotype well-suited for detecting gene knockouts that enhance RNAi activity. Based on our findings from this initial screen, we chose to use 6-day 100 μM 6TG treatment as the condition for CRISPR screen. To test the impacts of siRNA dosage on CRISPR screen, the actual CRISPR screen was done with both 150 nM GalNAc-*HPRT1* siRNA (low dose group), and 750 nM GalNAc-*HPRT1* siRNA (high dose group). The CRISPR screen experimental scheme is diagramed in Figure 2a. The genomic DNA samples were extracted from all sample pellets collected during the screen and sent to Cellecta for NGS (next gen sequencing) barcode sequencing.

**Figure 2.**
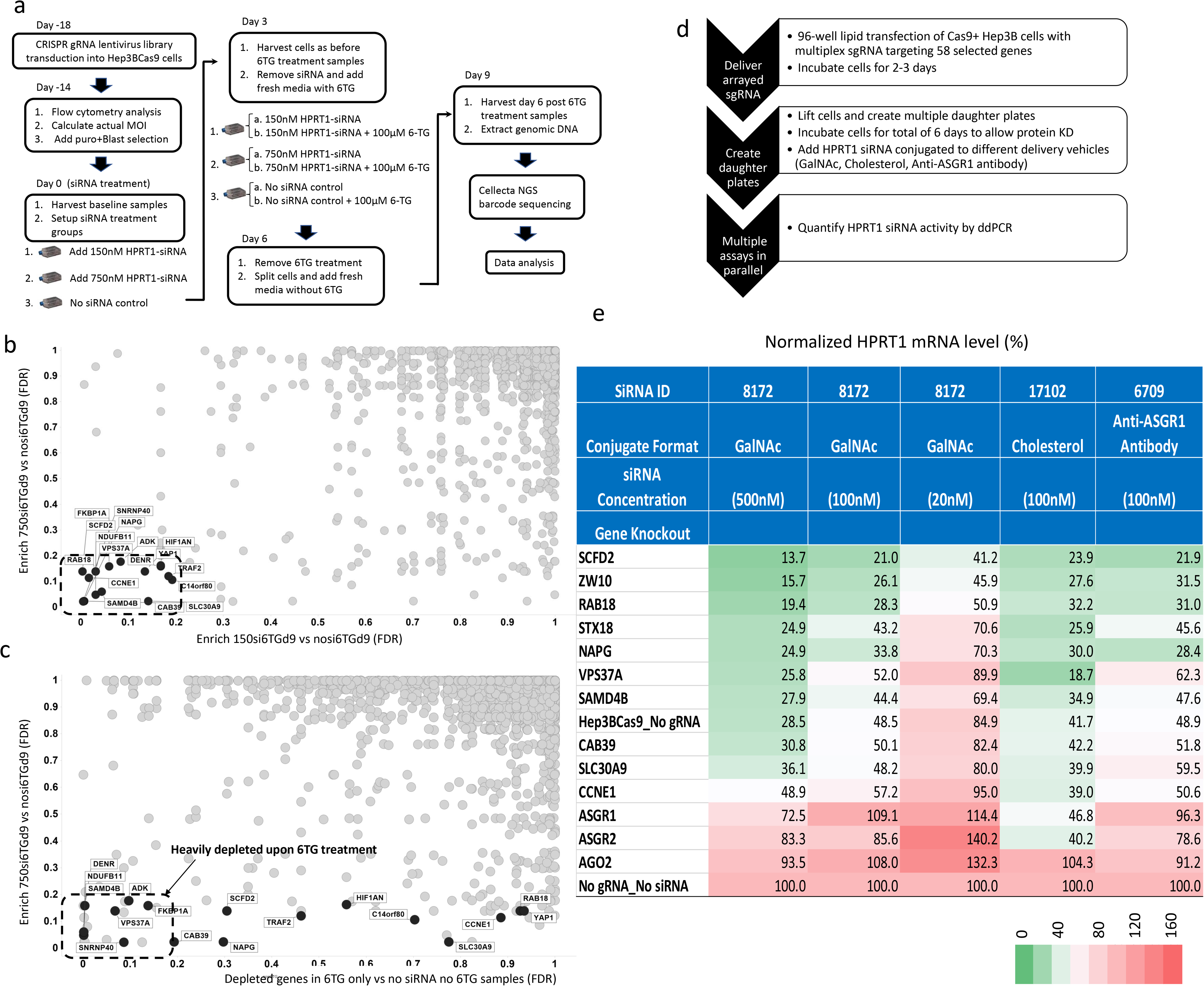
Large-scale pooled genome wide CRISPR knockout screen experiment and candidate gene validation. a) Experiment scheme of large-scale pooled genome wide CRISPR knockout screen. b) Analysis of the CRISPR screen results by overlapping enriched genes in both 150 nM siRNA + 6TG treated samples (150si6TGd9) vs. no siRNA but 6TG treated samples (nosi6TGd9) and 750 nM siRNA + 6TG treated samples (750si6TGd9) vs. no siRNA but 6TG treated samples (nosi6TGd9). A total of 17 genes were identified with FDR<0.2 (outlined by dashed line). c) Analysis of the CRISPR screen results by overlapping enriched genes from 750 nM siRNA + 6TG treated samples (750si6TGd9) vs. no siRNA but 6TG treated samples (nosi6TGd9) with depleted genes in 6TG only vs no siRNA no 6TG samples. The horizontal axis indicates the sensitivity to 6TG. The dashed line outlines 8 genes with FDR<0.2 that were heavily depleted upon 6TG treatment. d) Experiment scheme for testing of regulators of *HPRT1*-siRNA activity using secondary arrayed multiplexed synthetic gRNA screening in 96-well format. e) Heatmap results of secondary arrayed multiplexed synthetic gRNA shown in c). In the heatmap, red indicates reduced *HPRT1* siRNA silencing activity and green indicates enhanced *HPRT1* siRNA silencing activity. Colors indicate the percentage of *HPRT1/TBP* mRNA signals detected through ddPCR and normalized to no siRNA control. (*TBP*: a housekeeping gene)

### NGS sequencing results

The NGS sequencing results were analyzed by OGA algorithm^27^. False discovery rate (FDR) <0.2 was used as cutoff line. As shown in Supplementary Figure 3a-3b, all samples maintained good representation of gRNA library – roughly 77,000 gRNAs present with similar overall distribution. In addition, gRNAs that target *HPRT1* were successfully enriched by about 2-fold in 6TG treated vs. no 6TG group (Supplementary Figure 4). We then looked for additional genes that may play key roles in regulating GalNAc conjugated siRNA activity.

In order to identify genes that when knocked out can improve GalNAc conjugated siRNA internalization, trafficking or RNAi activity, we focused on gRNAs that were enriched in samples treated with both siRNA and 6TG but were not enriched in the 6TG treated only control group. These hits include genes that when knocked out could: 1) enhance GalNAc-*HPRT1* siRNA silencing potency, 2) increase sensitivity to 6TG in the absence of siRNA, or 3) enhance cell viability in the presence of 6TG. To select the genes with the most potent effects, we selected gene hits that were significantly (FDR<0.2) enriched in both high dose (750 nM) and low dose (150 nM) GalNAc-*HPRT1* siRNA and 6TG treated groups. (Figure 2b). This analysis identified 17 genes (Figure 2b). To understand whether any of these 17 genes have impacts on cells’ sensitivity to 6TG treatment in the absence of siRNA, we plotted these genes with the genes depleted in no siRNA but only 6TG treated group vs. no siRNA and no 6TG treated samples (Figure 2c). In Figure 2c, the horizontal axis indicates the sensitivity to 6TG. The genes that when knocked out enhance the sensitivity to 6TG and lead to strong cell death upon 6TG treatment are enriched on the horizontal axis with smaller FDR. If FDR < 0.2 was set as the cutoff, 8 genes were identified as promoting sensitivity to 6TG treatment (Figure 2c). The genes that when knocked out have no impact on 6TG sensitivity have larger FDR on horizontal axis, and these genes (*RAB18, YAP1, CCNE1, SLC30A9, C14orf80, HIF1AN, TRAF2, NAPG* and *SCFD2*) are the most interesting to us because their enrichment is most likely to be directly related to siRNA delivery and activity.

### Validation of primary CRISPR screen hits by secondary arrayed CRISPR screen

As discussed above, the HPRT1-6TG selection CRISPR screen may introduce false positive hits. To overcome this, we used a multiplexed synthetic gRNA system developed by Synthego (Redwood City, CA), which provides efficient knockout of target genes and can be scaled up to screen in 96-well or 384-well format without clonal isolation. In this multi-guide strategy, three gRNAs designed in close proximity to one another are delivered together to Cas9+ cells to induce a large deletion in the target gene and more efficient target gene knockout than individual gRNA. After inhouse validation of this strategy (Supplementary Figure 5), we ordered multiplexed synthetic gRNAs for some of the CRISPR screen hits along with control genes to run secondary arrayed CRISPR validation.

As illustrated in Figure 2d, the multiplexed synthetic gRNAs for genes identified in our initial CRISPR screen (*RAB18, CCNE1, SLC30A9, NAPG, SCFD2, VPS37A, SAMD4B* and *CAB39*) along with some control genes (*AGO2, ASGR1,* and *ASGR2*) were transfected into Hep3BCas9 cells. CRISPR-KO cells generated in this manner were then treated with GalNAc-*HPRT1* siRNA or *HPRT1* siRNA delivered through other conjugation formats (anti-ASGR1 antibody conjugated *HPRT1* siRNA (6709) and cholesterol conjugated *HPRT1* siRNA (17102), Supplementary Table 2). A heatmap of *HPRT1* siRNA silencing efficacy as measured by ddPCR (normalized to no siRNA control) is shown in Figure 2e. As expected, when *AGO2* is knocked out by multiplexed synthetic gRNA, the *HPRT1* siRNA silencing activity is abolished in all tested siRNA conjugates. Because ASGR1 is a critical component of ASGPR receptor, ASGR1 CRISPR-KO leads to loss of response to GalNAc-*HPRT1* siRNA as well as to anti-ASGR1 antibody conjugated *HPRT1* siRNA. However, knocking out *ASGR1* had no impacts on the function of Cholesterol conjugated *HPRT1* siRNA. These results indicate that the multiplexed synthetic gRNA system was working as expected. Some CRISPR screen hits: *RAB18, SCFD2, NAPG,* and *SAMD48* when knocked out by multiplexed synthetic gRNA enhanced siRNA effects to different degrees (Figure 2e). *VPS37A* specifically enhanced cholesterol conjugated siRNA efficacy. Other screen hits, *CAB39, CCNE1* and *SLC30A9,* could not be validated by the multiplexed synthetic gRNA approach. Proteins encoded by *ZW10* and *STX18* had been shown to interact with RAB18 protein^28,29^. Knocking out *ZW10* and *STX18* by multiplexed synthetic gRNA also enhanced siRNA silencing efficacy (Figure 2e).

### *RAB18* knockdown/knockout enhances the silencing effects of multiple siRNA conjugates

Since *RAB18* was the only RAB family member detected in our CRISPR screen, and because the RAB family is important in regulating intracellular vesicle trafficking, we decided to focus on understanding the mechanisms by which *RAB18* regulates siRNA activity in Hep3B. To study the function of *RAB18*, three *RAB18* specific siRNA molecules (siRAB18_1, siRAB18_2, and siRAB18_3) (Supplementary Table 4) purchased from Ambion were validated for their silencing potency of *RAB18* in Hep3B cells through transfection study. Among three tested siRNA molecules, siRAB18_3 which showed the best knocking down potency of *RAB18* (Figure 3a) was then used to study the function of *RAB18*. The Hep3B cells transfected with either siRAB18_3 or a non-targeting control siRNA molecule (siNTC) for 24hr were further treated with GalNAc-*HPRT1* siRNA at various concentration. As illustrated in Figure 3b, the siRNA18_3 treated cells were able to maintain low level (23.2%) of *RAB18* mRNA measured by ddPCR on day 4 post GalNAc-*HPRT1* siRNA treatment compared with siNTC treated cells. The level of *HPRT1* mRNA was also measured by ddPCR on day 4 post GalNAc-*HPRT1* siRNA treatment. As shown in Figure 3c, the knockdown of *HPRT1* was greater in siRAB18_3 treated Hep3B cells compared to siNTC treated Hep3B cells. The IC50 for siRAB18_3 treated cells or siNTC treated cells was 24.8nM versus 223.6nM (Figure 3c), respectively, a 10-fold change.

**Figure 3.**
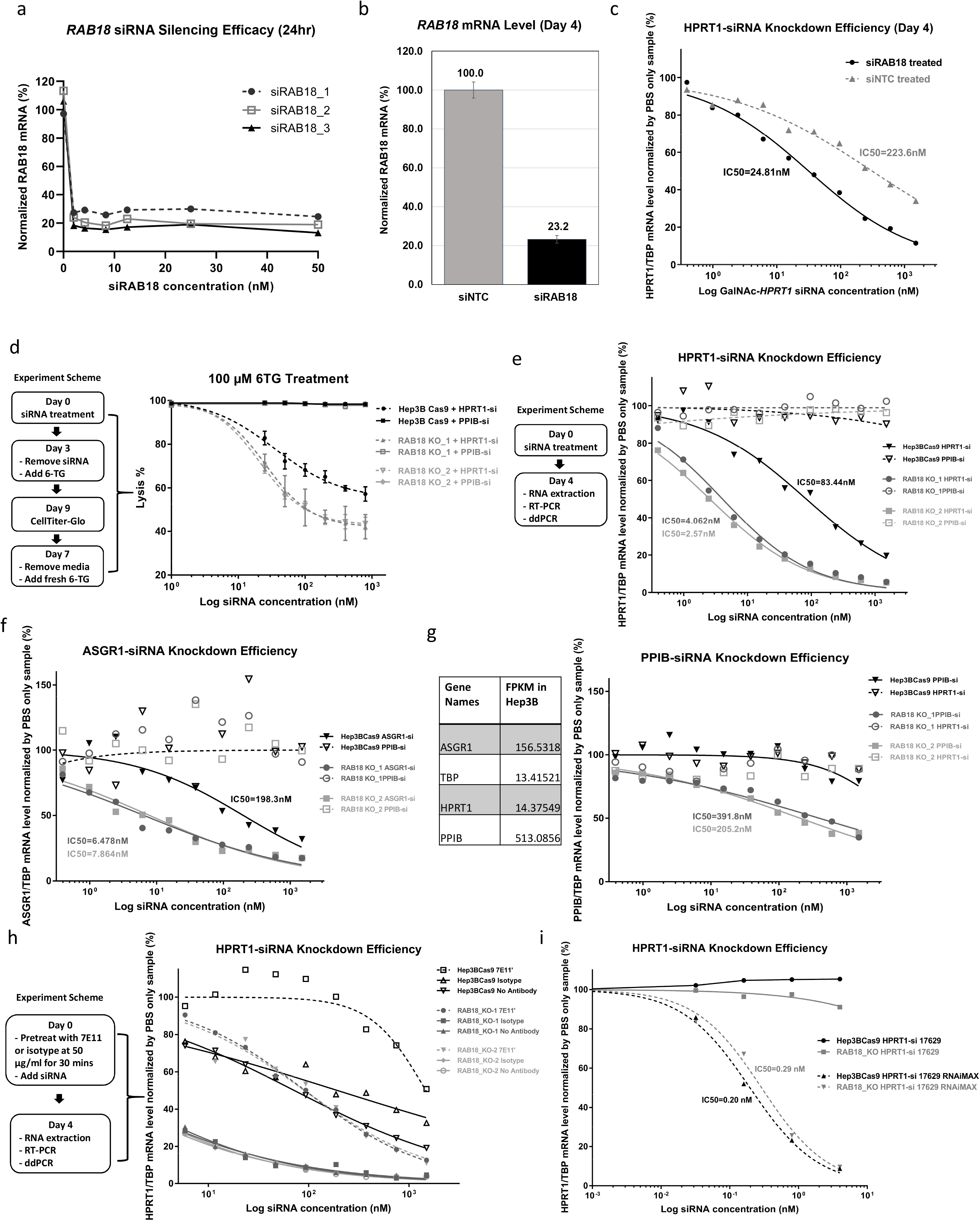
Validation of the effect of *RAB18* knockdown and knockout on siRNA silencing potency. a) The knocking down efficacy of three siRNA molecules targeting *RAB18* in Hep3B cells. The RNA samples were extracted from Hep3B cells treated with three different siRNA molecules targeting *RAB18* gene at various concentrations at 24hr post treatment. The cDNA samples synthesized from RNA through reverse transcription were then used to perform ddPCR. The *RAB18* ddPCR readings normalized by housekeeping gene *TBP* were used to calculate the percentage of *RAB18* mRNA level for this plot. b) The *RAB18* mRNA level measured by ddPCR on day 4 post GalNAc-*HPRT1* siRNA treatment. The Hep3B cells were pretreated with siRAB18_3 or siNTC molecules through transfection. 24hr later, after washing off the transfection media, the cells were treated with GalNAc-*HPRT1* siRNA at various concentrations. On day 4 post GalNAc-*HPRT1* siRNA treatment, the cells were harvested for ddPCR measurement of *RAB18* mRNA level. c) Measurement of GalNAc-*HPRT1* siRNA silencing potency in both siRAB18_3 and siNTC treated Hep3B cells by ddPCR on day 4 post-GalNAc-*HPRT1* siRNA treatment. The cells harvested from experiment described in “b” were also measured for *HPRT1* level through ddPCR. Plotted here are *HPRT1* mRNA levels in percentage normalized by housekeeping gene *TBP* readings and no siRNA (PBS only) treated control group. d) HPRT1-6TG live/dead selection performed in both Hep3BCas9 parental cells and *RAB18* knockout cells. Left panel outlines the experiment scheme and the right panel shows the cell lysis rate measured with CellTiter-Glo reagents (Promega, Madison, WI). e) Measurement of GalNAc-*HPRT1* siRNA silencing potency in both Hep3BCas9 and *RAB18* knockout cells by ddPCR on day 4 post-siRNA treatment. Left panel summarizes the experiment scheme and the right panel shows the *HPRT1* mRNA levels in percentage normalized to housekeeping gene *TBP* readings and no siRNA (PBS only) treated control group. *PPIB* siRNA was used as control siRNA. f) Measurement of GalNAc-*ASGR1* siRNA silencing potency in both Hep3BCas9 and *RAB18* knockout cells by ddPCR on day 4 post-siRNA treatment. The experiment scheme is the same as shown in “e”. Plotted here are *ASGR1* mRNA levels in percentage normalized by housekeeping gene *TBP* readings and no siRNA (PBS only) treated control group. *PPIB* siRNA was used as control siRNA. g) The same experiment shown in “e” was performed using GalNAc-*PPIB* siRNA. Left panel lists the normal target genes’ expression profile in FPKM (Fragments Per Kilobase of transcript per Million mapped reads) (obtained from Broad Institute Cancer Cell Line Encyclopedia (CCLE)). Right panel plots the *PPIB* mRNA levels in percentage normalized by housekeeping gene *TBP* readings and no siRNA (PBS only) treated control group. *HPRT1* siRNA was used as control siRNA. h) Antibody blocking test in Hep3BCas9 and *RAB18* knockout cells by using anti-ASGR1 antibody, 7E11. Left panel summarizes the experiment scheme and the right panel shows the ddPCR measurement of *HPRT1* mRNA levels in percentage normalized by housekeeping gene *TBP* readings and no siRNA (PBS only) treated control group. i) Unconjugated *HPRT1* siRNA transfection assay in Hep3BCas9 and *RAB18* knockout cells. The unconjugated *HPRT1* siRNA (17629) was delivered into Hep3BCas9 and *RAB18* knockout cells through either conjugating to GalNAc or Lipofectamine reagent (RNAiMAX) mediated transfection. The *HPRT1* mRNA levels were measured by ddPCR on day 4 after siRNA treatment. Plotted is the *HPRT1* mRNA levels in percentage normalized by housekeeping gene *TBP* readings and no siRNA (PBS only) treated control group. The features of siRNA 17629 is described in Supplementary Table 2.

In order to completely abolish the function of *RAB18*, we created two *RAB18* knockout pools (RAB18_KO_1 and RAB18_KO_2) by transducing two lentiviral gRNA vectors targeting *RAB18* (SIGMA vector: U6-gRNA: PGK-puro-2A-tagBFP) into Hep3BCas9 cells (Supplementary Figure 6a). The *RAB18* knockout efficiency was verified by Amplicon-EZ sequencing (GENEWIZ, Newbury Park, CA) (Supplementary Figure 6b and 6c). Knocking out *RAB18* did not alter cell viability in Hep3BCas9 cells (supplementary Figure 6d). Since *RAB18* was identified through HPRT1-6TG selection screen, we first repeated the same assay in *RAB18* knockout cells. As shown in Figure 3d, compared with the parental Hep3BCas9 cells, when treated with GalNAc-*HPRT1* siRNA approximately 15% more *RAB18* knockout cells were able to survive under 6TG selection (58% in *RAB18* knockout cells compared to 43% in Hep3BCas9 cells at the highest siRNA dose tested), indicating that *HPRT1* siRNA induced greater gene silencing in *RAB18* knockout cells than in Hep3BCas9 parental cells. Neither Hep3BCas9 cells nor *RAB18* knockout cells treated with GalNAc conjugated siRNA targeting *PPIB* gene (8714) as a non-relevant siRNA control showed enhanced resistance to 6TG treatment (Figure 3d, Supplementary Table 2). We then used ddPCR to directly measure the siRNA silencing potency in *RAB18* knockout cells. As illustrated in Figure 3e, 3c and 3d, Hep3BCas9 cells and *RAB18* knockout cells were treated with three GalNAc conjugated siRNAs: *HPRT1* siRNA, *ASGR1* siRNA (16084) (Supplementary Table 2), and *PPIB* siRNA. For all three tested siRNAs, the target gene knockdown was greater in *RAB18* knockout cells compared to Hep3BCas9 parental cells (Figure 3e-g). The IC50 for *HPRT1* siRNA in Hep3BCas9 or two *RAB18* knockout lines was 83.4nM versus 2.6nM or 4.1nM (Figure 3e), respectively, a 20~30-fold change. When tested using GalNAc-*ASGR1* siRNA, the IC50 was 198.3nM in Hep3BCas9 cells and 7.9nM or 6.5nM in two *RAB18* knockout cells (Figure 3f). Compared to *HPRT1* and *ASGR1*, *PPIB* is a highly abundantly expressed gene in Hep3B cells (Figure 3g), that could not be silenced by GalNAc-*PPIB* siRNA in Hep3BCas9 cells (Figure 3g). However, the same *PPIB* siRNA was able to silence *PPIB* in two *RAB18* knockout pools (IC50=205.2nM or 391.8nM) (Figure 3g). The siRNA silencing efficacy at a later time point (11 days) was also checked (Supplementary Figure 7a-c). Although the silencing effect declined as the cells proliferated over time, the silencing potency was greater in *RAB18* knockout cells than in Hep3BCas9 cells. For example, when treated with GalNAc-*ASGR1*, the IC50 at day 11 was 363.6nM in Hep3BCas9 cells and 41.3nM or 58.3nM in two *RAB18* knockout pools (Supplementary Figure 7a). These results lead us to conclude that *RAB18* knockout enhances the silencing potency of GalNAc conjugated siRNA as well as cholesterol and antibody conjugated siRNA.

### Gene silencing induced by GalNAc conjugated siRNA in *RAB18* knockout cells requires ASGR1

As discussed and tested earlier, the GalNAc siRNA conjugate induced gene silencing is mediated through ASGR1. We therefore tested if ASGR1 was required for GalNAc siRNA conjugates to function in *RAB18* knockout cells using an antibody blocking test (Figure 3h, Supplementary Figure 7d-e). As shown in Figure 3h, the application of 7E11 was able to reduce the siRNA silencing efficacy of *HPRT1* gene in Hep3BCas9 and *RAB18* knockout cells. Similar results were obtained when the same experiment performed by using *ASGR1* siRNA and *PPIB* siRNA to silence *ASGR1* and *PPIB* (Supplementary Figure 7d-e). Similarly to what we observed in Hep3B cells, the GalNAc siRNA conjugates rely on ASGR1 to enter *RAB18* knockout cells. The two individually generated *RAB18* knockout pools behaved identically in all tests. Therefore, only one *RAB18* knockout pool was used for the rest of the related experiments (referred as RAB18_KO).

### *RAB18* knockout shows no effect on the activity of siRNA delivered through Lipofectamine transfection

Lipofectamine reagents have been widely used experimentally as a safe and efficient method to deliver exogenous DNA and RNA into cells. After confirming that knocking out *RAB18* enhances siRNA potency delivered through GalNAc conjugates, we asked if knocking out *RAB18* could enhance siRNA potency delivered through Lipofectamine mediated transfection. To address this question, we used an unconjugated *HPRT1* siRNA (17629) (Supplementary Table 2) to treat the parental Hep3BCas9 and *RAB18* knockout cells at various concentrations with or without Lipofectamine RNAiMAX reagent (Invitrogen, Waltham, MA). As summarized in Figure 3i, Lipofectamine reagents efficiently silenced the target gene *HPRT1* at similar levels in both Hep3BCas9 (IC50=0.2nM) and *RAB18* knockout cells (IC50=0.3nM). This finding indicates that *RAB18* does not alter Lipofectamine mediated siRNA activity.

## DISCUSSION

The HPRT1-6TG selection based CRISPR-Cas9 screen performed in Hep3B background has successfully identified several key regulators of GalNAc conjugated siRNA activity. Some of the hits from this screen, such as *RAB18, SCFD2, NAPG,* and *VPS37A,* have been validated through a secondary arrayed CRISPR screen system by using multiplexed synthetic gRNA. Here, we focused our efforts on studying the effects *RAB18* on siRNA activity.

Having confirmed that knocking out *RAB18* enhances siRNA silencing potency on multiple tested target genes (*HPRT1, ASGR1,* and *PPIB*) and through multiple siRNA conjugated formats (GalNAc, Cholesterol, and antibody conjugates), we attempted to elucidate the functional linkage between *RAB18* and siRNA activity. RAB GTPases constitute the largest family of small GTPases that have important roles in regulating membrane trafficking through switching between GTP-bound ‘on’ form and GDP-bound ‘off’ forms. There are more than 60 RAB family members in humans that are localized to distinct intracellular membranes and play important roles in regulating intracellular vesicle budding, uncoating, motility, and fusion. Once internalized, siRNA has been shown to traffic through the endocytic pathway^30,31^. We therefore expected multiple members of the RAB family to be identified in our CRISPR screen as regulators of siRNA activity. To our surprise, *RAB18* was the only RAB family member that came out of our screen (FDR<0.2).

As one of the 20 most highly conserved RAB GTPases present in the last eukaryotic common ancestor of both the plant and animal kingdoms^32,33^, RAB18 has attracted great research interest and attention. Studies conducted in the last couple decades have linked *RAB18* to regulation of lipid droplet (LD) formation^28,34^, inhibition of COPI independent retrograde trafficking from Golgi to endoplasmic reticulum (ER)^35^, regulation of secretory granules^36^ and peroxisomes^37^, promotion of hepatitis C virus (HCV) assembly on the LD membrane^38^, and regulation of normal ER structure^39^. Despite the intensive efforts on studying *RAB18*, a defined molecular function of *RAB18* and its site of action has remained elusive. It is very difficult to tease out what known functions of *RAB18* gene might contribute to regulation of siRNA activity, or whether a novel function of *RAB18* needs to be identified. However, several lines of evidence may provide a clue to a potential mechanism. First, the NRZ (NAG-RINT1-ZW10) tethering factors and their associated ER-localized SNAREs (Use1, Syntaxin18, and BNIP1) form a complex with GTP-bound form of RAB18 protein to mediate ER-LD contact formation^28,29^. Knocking out the genes *ZW10* and *STX18* (encoding Syntaxin18) by multiplexed synthetic gRNA enhanced siRNA silencing efficacy (Figure 2e). This indicates that genes interacting with *RAB18* to regulate ER-LD tethering have the same function in inhibiting siRNA silencing activity. Although it is still not clear how the ER-LD tethering impacts siRNA efficacy, this finding guided our attention to ER. We then asked, what is the connection between ER and siRNA silencing? The siRNA mediated degradation of target mRNA has been shown to take place in the cytoplasm^40^. However, the subcellular sites of RNA silencing remain under debate. Intriguingly ER as a site for protein translation mediated by ribosomes has been shown to be a central nucleation site of siRNA mediated RNA silencing^41^. In addition, an ER membrane resident protein CLIMP-63 has been proven to interact with and stabilize Dicer^42^. As indicated in these studies, ER might serve as a subcellular silencing site for siRNA. After being internalized into endosomes, the siRNA inside endosomes could travel to ER through retrograde transport. There are two different pathways of retrograde transport: the COPI-dependent and the COPI-independent pathways. Interestingly, *RAB18* loss of function mutants had been shown to specifically enhance COPI-independent retrograde Golgi-ER transport^35^. Although the exact molecular mechanism of *RAB18* regulation of siRNA activity has not yet been elucidated, the fact that knocking out genes functioning together with *RAB18* in regulating LD and ER tethering, such as *ZW10* and *STX18*, has similar impacts on siRNA silencing potency (Figure 2e) suggests that ER and retrograde transport related regulation might be worth more attention.

Despite the success of our HPRT1-6TG selection screen in identifying candidate regulators of siRNA potency, there were some limitations to this approach. First, the live/dead selection represents a very harsh cutoff for improving siRNA silencing potency. The idea behind HPRT-6TG selection is to use siRNA to knockdown *HPRT1* gene. Resistance to 6TG is associated with the extent of knockdown of *HPRT1*^43^. Our follow up study of *RAB18* has shown that by knocking out *RAB18*, the siRNA IC50 dose could be reduced by 20 to 30-fold (Figure 3e), indicating that *RAB18* is a strong regulator of siRNA activity. Nevertheless, when both Hep3BCas9 and *RAB18* knockout cells were challenged with HPRT1-6TG selection, the cell survival rate was only changed from 43% in Hep3BCas9 cells to 58% in *RAB18* knockout cells at the highest siRNA dosage tested (Figure 3d). This suggests that gene knockouts that improve survival by less than this amount may not be detected, even if they improve siRNA efficacy. Second, genes that regulate the sensitivity to 6TG could also be identified from the screen without having any function related to siRNA activity. These could include genes such as *HPRT1* itself as well as genes involved in the mismatch repair pathway^26^. Third, siRNA induced gene silencing is a complex process, in which multiple genes may be required to regulate individual steps. Therefore, one gRNA per cell strategy will miss the redundant genes. Finally, the screen described here is focused toward identifying genes that confer resistance but not genes that sensitize the siRNA activity. Knocking out genes that are normally required for siRNA activity would lead to resistance to siRNA activity and cause the cells to be depleted upon 6TG selection; consequently their corresponding gRNAs would be depleted from NGS sequencing along with other gRNAs that cause cell death and would not be detected by our approach.

The described CRISPR screen was performed in Hep3B cells. Despite the great success of delivering siRNA to liver through conjugating to GalNAc, delivering siRNA into other tissues is still challenging. While it is still unclear whether the same siRNA trafficking route is utilized in other tissue or cell types, *RAB18* is a universally expressed gene across multiple tissue types and is highly conserved across species. It would therefore be interesting to see if *RAB18* knockout in other cell or tissue types can also enhance siRNA activity.

We report here the identification, using a pooled genome-wide CRISPR-Cas9 screen, of a single gene (*RAB18*) that, when knocked out, can enhance siRNA mediated gene silencing by at least 20-fold (IC50) in Hep3B cells. Given the current interest in utilizing siRNA as a therapeutic modality, identification of this key regulator may allow for the development of future pharmacological strategies to enhance siRNA efficacy.

## MATERIALS AND METHODS

### Cell line and culture condition

The Hep3B cells were purchased from ATCC (Manassas, VA). The culture condition for Hep3B cells is: EMEM (Eagle’s Minimum Essential Medium from ATCC, Cat# 30-2003) + 10% FBS (Fetal Bovine Serum). The culture condition for Hep3Bcas9 cells is: EMEM + 10% FBS + 10 μg/mL Blasticidin. And the culture condition for *RAB18* knockout cells is: EMEM + 10% FBS + 10 μg/mL Blasticidin + 0.5 μg/mL Puromycin.

### Generate Hep3BCas9 cells and validate their editing function

A TransEDIT CRISPR Cas9 nuclease expression lentivirus (pCLIP-Cas9-Nuclease-EFS-Blast) ordered from TransOMIC technologies (Huntsville, AL, Cat# NC0956087) was transduced at multiple MOI (0.5, 1, and 2) into Hep3B cells to generate Cas9 stable pools: Hep3BCas9_0.5, Hep3BCas9_1, and Hep3BCas9_2, respectively. All cells were selected and maintained with 10 μg/mL Blasticidin after transduction. No toxicities were observed in all Cas9 stable expression Hep3B pools. Two gRNA lentivirus vectors targeting *SLC3A2* and *ASGR1* ordered from Milipore Sigma (Supplementary Table 3) were transduced individually into both parental Hep3B cell line and each of the Cas9 stable Hep3B pool. The *SLC3A2* and *ASGR1* expression levels before and after gRNA lentivirus transduction were measured through antibody staining followed by FLOW cytometry analysis. Compared to the parental Hep3B cell line, both target genes were successfully knocked out in all Cas9 stable Hep3B pools (Supplementary Figure 1), demonstrated the Cas9 stable Hep3BCas9 cells were fully equipped with editing function. Since the editing effects were similar in all three Cas9 stable Hep3B pools, the one with lowest MOI (0.5, referred as Hep3BCas9 in the rest of this article) was chosen to perform the CRISPR screen to search for key regulators of GalNAc conjugated siRNA induced silencing to minimized potential Cas9 toxicity.

### HPRT1-6TG selection test

To avoid both insufficient and over killing caused by 6TG, the feasibility of using HPRT1-6TG live/dead selection for CRISPR screen was tested in a small-scale pilot run using 100 μM 6TG (a dose close to IC70, Figure 1d) and 20 μM 6TG (a dose close to IC50, Supplementary Figure 2). The cells were first equally divided into four groups (0.6E+06 cells/group): 1) siRNA only, 2) siRNA with 6TG treatment, 3) 6TG only, and 4) negative control. To obtain sufficient but not excessive siRNA effect, a 750nM (about IC60) GalNAc-*HPRT1* siRNA (8172) was added to group 1 and 2 on day 0 of experiment. On day 3 of experiment, the tissue culture media was removed from each group and then 100 μM 6TG (or 20 μM 6TG) was added to group 2 and 3, while non-selection full growth media was added to group 1 and 4. The cells were incubated for 3 days after 6TG treatment. Then cells were then split and the 6TG media was replaced with full growth media without 6TG and cultured for additional 3 days. The cell count readings (measured by ViCell) were recorded on day 3 post-6TG treatment and day 6 post-6TG treatment and plotted in Figure 1d and Supplementary Figure 2.

### Large scale pooled genome wide CRISPR screen

An 80k genome wide CRISPR gRNA lentivirus library (CRISPR KOHGW 80K (lot# 17050301)) was purchased from Cellecta (Mountain View, CA) to generate a gene knockout pool. The CRISPR KOHGW 80K library is constructed in Cellecta’s pRSG16-U6-sg-UbiC-TagRFP-2A-Puro lentiviral vector that expresses gRNA under a wild-type U6 promoter and TagRFP and Puro resistance genes under a human ubiquitin C promoter. This library covers approximately 19,000 genes with 4 gRNA for each gene. The procedure of large scale CRISPR screen is illustrated in Figure 2a. Briefly, the gRNA lentivirus library was transduced into 9.2E+07 Hep3BCas9 cells. The actual library transduction efficiency as reflected by RFP positive cell population (61%) was checked through flow cytometry analysis on day 4 post-transduction. Based on calculation, the actual gRNA lentivirus library transduction MOI was about 0.9, and the actual coverage was 1035. The transduced cells were then selected with puromycin and blasticidin for 14 days. On day 14 post-selection, 87% of the cells were RFP positive (indicating 87% of the cells had an integrated gRNA) by flow cytometry. On day 14 post-selection, 1E+08 cells were collected and frozen as baseline sample. The rest of cells were equally divided into three groups (2.4E+08 cells/group): group 1 was treated with 150 nM GalNAc-*HPRT1* siRNA as low dose group, group 2 was treated with 750 nM GalNAc-*HPRT1* siRNA as high dose group, and group 3 was set as no siRNA control. On day 3 post-siRNA treatment, 2E+08 cells were collected and frozen from each group as before 6TG treatment samples, then the rest of the cells in each group were further divided into two subgroups: a) no 6TG group and b) 6TG group. The cell culture medium with siRNA was removed from each flask and fresh medium containing 100 μM 6TG was added into each flask of 6TG groups and fresh medium without 6TG was added to each flask in no 6TG groups. All cells were incubated for another 3 days then all cells were split into fresh medium without 6TG. After a final 3-day incubation, all cells were harvested. The genomic DNA samples were extracted from all samples collected by using Gentra Puregene Cell Kit (QIAGEN INC, Cat# 158767) following the user manual and sent to Cellecta for NGS barcode sequencing.

### Secondary arrayed CRISPR screen

The multiplexed synthesized gRNA of each target gene for secondary arrayed CRISPR screen was designed and synthesized by Synthego Corporation (Palo Alto, CA). All gRNAs were transfected into Hep3BCas9 stable cells at 96-well plate format using Lipofectamine CRISPRMAX Cas9 Transfection Reagent (Invitrogen, Cat# CMAX00008). 1.5 μL of 0.3 uM multiplexed synthesized gRNA was first mixed with 8.5 μL Opti-MEM medium in each well. 0.2 μL of CRISPRMAX reagent diluted in 5 μL of Opti-MEM medium was then added to each well and incubated at room temperature for 5 to 10 minutes. After incubation, 85 μL (15,000 cells per well) of Hep3BCas9 stable cells were added to each well. The plate was allowed to sit for 20 minutes prior to placing it in 37°C tissue culture incubator, and transfection medium was replaced with EMEM containing 10% FBS and 1% AA (Antibiotic Antimycotic Solution) at ~6 hours after transfection. The cells were split at 1:6 ratio on day 3 post-incubation. The cells were incubated for a total of 6 days after CRISPRMAX transfection to allow protein knockdown. On day 6 post-transfection, *HPRT1* siRNA conjugated to different delivery vehicles (GalNAc, Cholesterol, Anti-ASGR1 antibody) was added to each well at the desired concentrations (500 nM, 100 nM and 20 nM) followed by 4-day incubation period in 37°C tissue culture incubator. The total RNA of each sample was extracted by using KingFisher Flex System (Thermo Fisher Scientific) and MagMAX mirVana Total RNA Isolation Kit (Applied Biosystems, Cat# A27828) as per manufacturer instructions. The cDNA was then synthesized from total RNA sample using the Applied Biosystems High Capacity Reverse Transcription Kit (Cat# 4368813), and used to quantify siRNA activity by ddPCR (Droplet Digital Polymerase Chain Reaction).

### Droplet Digital Polymerase Chain Reaction (ddPCR)

The ddPCR reactions were assembled using BioRad’s ddPCR Supermix for Probes (Cat# 1863010) as per the user manual. Droplets were then generated by QX200 Automated Droplet Generator (BioRad, Cat# 1864101). Thermal cycling reactions were then performed on C1000 Touch Thermal Cycler with 96-Deep Well Reaction Module (BioRad, Cat# 1851197) (BioRad, Cat# 1851197). The reactions were then read by QX200 Droplet Reader (BioRad, Cat# 1864003) and analyzed by using BioRad’s QuantaSoft software package. The predesigned primer/probe for ddPCR assays were obtained from Integrated DNA Technologies (Coralville, IA) with 3.6:1 primer to probe ratio. The assay ID of primer/probe used for quantifying *HPRT1* gene is: Hs.PT.39a.22214821. The assay ID of primer/probe used for quantifying *ASGR1* gene is: Hs.PT.56a.24725395. The assay ID of primer/probe used for quantifying *PPIB* gene is: Hs.PT.58.40006718. The assay ID of primer/probe used for quantifying housekeeping gene *TBP* is: Hs.PT.58.19489510. The ddPCR copy number readings (copies/20 μL) of both target gene (*HPRT1, ASGR1* or *PPIB*) and housekeeping gene *TBP* were recorded for each well. The normalized target gene mRNA level was calculated by dividing the ddPCR reading of the target gene by the ddPCR reading of *TBP* taken from the same well. The resulting number of siRNA treated sample was further divided by the number of no siRNA treatment sample to obtain the percentage reading of the target gene mRNA level, which was plotted in Figure 1b, Figure 3e~i, and Figure 4.

### siRAB18 and siNTC transfection

The siRNA molecules targeting *RAB18* gene, siRAB18_1 (Ambion Silencer Select cat# 4390824 ID# s22703), siRAB18_2 (ID# s22704), and siRAB18_3 (ID# s22705) were purchased from Ambion (Ambion, Austin, TX). The non-targeting negative control siRNA (siNTC, cat# 4390843) were purchased from Invitrogen. The sequence details of siRNA targeting *RAB18* were described in Supplementary Table 4. To test siRAB18 efficacy, several concentrations of each siRAB18 molecule (0.24nM to 50nM) or sterile water (negative control) was individually reverse transfected in duplicate into Hep3B cells using lipofectamine RNAiMAX (Invitrogen, cat#13778075). 24 hours post-transfection, cells were lysed and harvested for RNA using MagMAX mirVana Total RNA Isolation kit (Applied Biosystems, Cat# A27828) and reverse transcribed for ddPCR analysis using the Applied Biosystems High Capacity Reverse Transcription Kit (Cat# 4368813), according to manufacturer instructions. For analysis of the effect of *RAB18* knockdown on GalNAc-*HPRT1* siRNA efficacy, siNTC (50nM) or siRAB18-3 (50nM) was reverse transfected into Hep3B cells. 24 hours post-transfection, cells were trypsinized and washed twice in EMEM to remove residual transfection reagent, then plated into 96-well plates containing either PBS or multiple concentrations of GalNAc-*HPRT1* siRNA. On day 4 post GalNAc-*HPRT1* siRNA treatment, the cells were lysed for RNA isolation and cDNA synthesis as described above.

### Anti-ASGR1 antibody blocking test

The Hep3BCas9 cells and *RAB18* knockout cells were first pre-incubated with in-house generated anti-ASGR1 antibody (7E11), isotype control antibody, or no antibody for half an hour, followed by adding GalNAc-*HPRT1* siRNA treatment at different doses. The final antibody concentration was 50 μg/mL and 2,000 cells were seeded each well. After incubating in 37°C tissue culture incubator for 4 days, the target gene (*HPRT1*) mRNA levels were measured using ddPCR analysis.

## Supporting information

All Supplementary Figures

## ACKNOWLEDGEMENTS

We thank Stephen Wong and Oliver Homann for providing input for NGS data analysis. We also thank Karen Siegler for providing 7E11 antibody.

## AUTHOR CONTRIBUTIONS

Michael Ollmann, Jiamiao Lu, Patrick Collins, Chi-Ming Li, and Songli Wang conceived and designed the study. Jiamiao Lu carried out the pooled genome wide CRISPR screen, ddPCR analysis, *RAB18* knockout study and drafted the manuscript. Elissa Swearingen carried out initial siRNA efficacy test and 6TG sensitivity test in Hep3B cells. Miki Hardy conducted the secondary arrayed CRISPR screen. Patrick Collins performed the statistical analysis of NGS results. Bin Wu conjugated siRNA molecules tested in this study. Eric Yuan performed *RAB18* knockdown study. Daniel Lu carried out Amplicon_Seq analysis to assess the editing efficacy of arrayed CRISPR platform. All authors contributed to manuscript revisions. All authors approved the final version of the manuscript and agree to be held at countable for the content therein.

## REFERENCES

1 Chu, C. Y. & Rana, T. M. Small RNAs: regulators and guardians of the genome. J Cell Physiol 213, 412–419, doi:10.1002/jcp.21230 (2007).

2 Dowdy, S. F. Overcoming cellular barriers for RNA therapeutics. Nat Biotechnol 35, 222–229, doi:10.1038/nbt.3802 (2017).

3 Juliano, R. L. The delivery of therapeutic oligonucleotides. Nucleic Acids Res 44, 6518–6548, doi:10.1093/nar/gkw236 (2016).

4 Khvorova, A. & Watts, J. K. The chemical evolution of oligonucleotide therapies of clinical utility. Nat Biotechnol 35, 238–248, doi:10.1038/nbt.3765 (2017).

5 Nair, J. K., Attarwala, H., Sehgal, A., Wang, Q., Aluri, K., Zhang, X., Gao, M., Liu, J., Indrakanti, R., Schofield, S., et al. Impact of enhanced metabolic stability on pharmacokinetics and pharmacodynamics of GalNAc-siRNA conjugates. Nucleic Acids Res 45, 10969–10977, doi:10.1093/nar/gkx818 (2017).

6 Nair, J. K., Willoughby, J. L., Chan, A., Charisse, K., Alam, M. R., Qang, Q., Hoekstra, M., Kandasamy, P., Kel’in, A. V., Milstein, S. et al. Multivalent N-acetylgalactosamine-conjugated siRNA localizes in hepatocytes and elicits robust RNAi-mediated gene silencing. J Am Chem Soc 136, 16958–16961, doi:10.1021/ja505986a (2014).

7 Chan, A., Liebow, A., Yasuda, M., Gan, L., Racie, T., Maier, M., Kuchimanchi, S., Foster, D., Milstein, S., Charisse, K. et al. Preclinical Development of a Subcutaneous ALAS1 RNAi Therapeutic for Treatment of Hepatic Porphyrias Using Circulating RNA Quantification. Mol Ther Nucleic Acids 4, e263, doi:10.1038/mtna.2015.36 (2015).

8 Pasi, K. J., Rangarajan, S., Georgiev, P., Mant, T., Creagh, M. D., Lissitchkov, T., Bevan, D., Austin, S., Hay, C. R., Hegemann, I. et al. Targeting of Antithrombin in Hemophilia A or B with RNAi Therapy. N Engl J Med 377, 819–828, doi:10.1056/NEJMoa1616569 (2017).

9 Baenziger, J. U. & Fiete, D. Galactose and N-acetylgalactosamine-specific endocytosis of glycopeptides by isolated rat hepatocytes. Cell 22, 611–620, doi:10.1016/0092-8674(80)90371-2 (1980).

10 Meier, M., Bider, M. D., Malashkevich, V. N., Spiess, M. & Burkhard, P. Crystal structure of the carbohydrate recognition domain of the H1 subunit of the asialoglycoprotein receptor. J Mol Biol 300, 857–865, doi:10.1006/jmbi.2000.3853 (2000).

11 Pricer, W. E., Jr. & Ashwell, G. The binding of desialylated glycoproteins by plasma membranes of rat liver. J Biol Chem 246, 4825–4833 (1971).

12 Spiess, M. & Lodish, H. F. An internal signal sequence: the asialoglycoprotein receptor membrane anchor. Cell 44, 177–185, doi:10.1016/0092-8674(86)90496-4 (1986).

13 Braun, J. R., Willnow, T. E., Ishibashi, S., Ashwell, G. & Herz, J. The major subunit of the asialoglycoprotein receptor is expressed on the hepatocellular surface in mice lacking the minor receptor subunit. J Biol Chem 271, 21160–21166, doi:10.1074/jbc.271.35.21160 (1996).

14 Drickamer, K., Mamon, J. F., Binns, G. & Leung, J. O. Primary structure of the rat liver asialoglycoprotein receptor. Structural evidence for multiple polypeptide species. J Biol Chem 259, 770–778 (1984).

15 Prakash, T. P., Graham, M. J., Yu, J., Carty, R., Low, A., Chappell, A., Schmidt, K., Zhao, C., Aghajan, M., Murray, H. F. et al. Targeted delivery of antisense oligonucleotides to hepatocytes using triantennary N-acetyl galactosamine improves potency 10-fold in mice. Nucleic Acids Res 42, 8796–8807, doi:10.1093/nar/gku531 (2014).

16 Gilleron, J., Querbes, W., Zeigerer, A., Borodovsky, A., Marsico, G., Schubert, U., Manygoats, K., Seifert, S., Andree, C., Stoter, M. et al. Image-based analysis of lipid nanoparticle-mediated siRNA delivery, intracellular trafficking and endosomal escape. Nat Biotechnol 31, 638–646, doi:10.1038/nbt.2612 (2013).

17 Leuschner, P. J., Ameres, S. L., Kueng, S. & Martinez, J. Cleavage of the siRNA passenger strand during RISC assembly in human cells. EMBO Rep 7, 314–320, doi:10.1038/sj.embor.7400637 (2006).

18 Nakanishi, K. Anatomy of RISC: how do small RNAs and chaperones activate Argonaute proteins? Wiley Interdiscip Rev RNA 7, 637–660, doi:10.1002/wrna.1356 (2016).

19 Springer, A. D. & Dowdy, S. F. GalNAc-siRNA Conjugates: Leading the Way for Delivery of RNAi Therapeutics. Nucleic Acid Ther 28, 109–118, doi:10.1089/nat.2018.0736 (2018).

20 Cho, S. W., Kim, S., Kim, J. M. & Kim, J. S. Targeted genome engineering in human cells with the Cas9 RNA-guided endonuclease. Nat Biotechnol 31, 230–232, doi:10.1038/nbt.2507 (2013).

21 Cong, L., Ran, F. A., Cox, D., Lin, S., Barretto, R., Habib, N., Hsu, P. D., Wu, X., Jiang, W., Marraffini, L. A. et al. Multiplex genome engineering using CRISPR/Cas systems. Science 339, 819–823, doi:10.1126/science.1231143 (2013).

22 Jinek, M., Chylinski, K., Fonfara, I., Hauer, M., Doudna, J. A., Charpentier, E. A programmable dual-RNA-guided DNA endonuclease in adaptive bacterial immunity. Science 337, 816–821, doi:10.1126/science.1225829 (2012).

23 Wang, T., Wei, J. J., Sabatini, D. M. & Lander, E. S. Genetic screens in human cells using the CRISPR-Cas9 system. Science 343, 80–84, doi:10.1126/science.1246981 (2014).

24 Shalem, O., Sanjana, N. E., Hartenian, E., Shi, X., Scott, D. A., Mikkelson, T., Heckl, D., Ebert, B. L., Root, D. E., Doench, J. G. et al. Genome-scale CRISPR-Cas9 knockout screening in human cells. Science 343, 84–87, doi:10.1126/science.1247005 (2014).

25 Liao, S., Tammaro, M. & Yan, H. Enriching CRISPR-Cas9 targeted cells by co-targeting the HPRT gene. Nucleic Acids Res 43, e134, doi:10.1093/nar/gkv675 (2015).

26 Glaab, W. E., Risinger, J. I., Umar, A., barrett, J. C., Kunkel, T. A., Tindall, K. R. Resistance to 6-thioguanine in mismatch repair-deficient human cancer cell lines correlates with an increase in induced mutations at the HPRT locus. Carcinogenesis 19, 1931–1937, doi:10.1093/carcin/19.11.1931 (1998).

27 Meisen, W. H., Nejad, Z. B., Hardy, M., Zhao, H., Oliverio, O., Wang, S., Hale, C., Ollmann, M. M., Collins, P. J. Pooled Screens Identify GPR108 and TM9SF2 as Host Cell Factors Critical for AAV Transduction. Mol Ther Methods Clin Dev 17, 601–611, doi:10.1016/j.omtm.2020.03.012 (2020).

28 Xu, D., Li, Y., Wu, L., Li, Y., Zhao, D., Yu, J., Huang, T., Ferguson, C., Parton, R. G., Yang, H. et al. Rab18 promotes lipid droplet (LD) growth by tethering the ER to LDs through SNARE and NRZ interactions. J Cell Biol 217, 975–995, doi:10.1083/jcb.201704184 (2018).

29 Li, D., Zhao, Y. G., Li, D., Zhao, H., Huang, J., Miao, G., Feng, D., Liu, P., Li, D., Zhang, H. The ER-Localized Protein DFCP1 Modulates ER-Lipid Droplet Contact Formation. Cell Rep 27, 343–358 e345, doi:10.1016/j.celrep.2019.03.025 (2019).

30 Pereira-Leal, J. B. & Seabra, M. C. Evolution of the Rab family of small GTP-binding proteins. J Mol Biol 313, 889–901, doi:10.1006/jmbi.2001.5072 (2001).

31 Stenmark, H. & Olkkonen, V. M. The Rab GTPase family. Genome Biol 2, REVIEWS3007, doi:10.1186/gb-2001-2-5-reviews3007 (2001).

32 Elias, M., Brighouse, A., Gabernet-Castello, C., Field, M. C. & Dacks, J. B. Sculpting the endomembrane system in deep time: high resolution phylogenetics of Rab GTPases. J Cell Sci 125, 2500–2508, doi:10.1242/jcs.101378 (2012).

33 Klopper, T. H., Kienle, N., Fasshauer, D. & Munro, S. Untangling the evolution of Rab G proteins: implications of a comprehensive genomic analysis. BMC Biol 10, 71, doi:10.1186/1741-7007-10-71 (2012).

34 Martin, S., Driessen, K., Nixon, S. J., Zerial, M. & Parton, R. G. Regulated localization of Rab18 to lipid droplets: effects of lipolytic stimulation and inhibition of lipid droplet catabolism. J Biol Chem 280, 42325–42335, doi:10.1074/jbc.M506651200 (2005).

35 Dejgaard, S. Y., Murshid, A., Erman, A., Kizilay, O., Verbich, D., Lodge, R., Dejgaard, K., Ly-Hartig, T. B., Pepperkok, R., Simpson, J. C. et al. Rab18 and Rab43 have key roles in ER-Golgi trafficking. J Cell Sci 121, 2768–2781, doi:10.1242/jcs.021808 (2008).

36 Vazquez-Martinez, R., Cruz-Garcia, D., Duran-Prado, M., Peinado, J. R., Castano, J. P., Malagon, M. M. Rab18 inhibits secretory activity in neuroendocrine cells by interacting with secretory granules. Traffic 8, 867–882, doi:10.1111/j.1600-0854.2007.00570.x (2007).

37 Gronemeyer, T., Wiese, S., Grinhagens, S., Schollenberger, L., Satyagraha, A., Huber, L. A., Meyer, H. E., Warscheid, B., Just, W. W. Localization of Rab proteins to peroxisomes: a proteomics and immunofluorescence study. FEBS Lett 587, 328–338, doi:10.1016/j.febslet.2012.12.025 (2013).

38 Salloum, S., Wang, H., Ferguson, C., Parton, R. G. & Tai, A. W. Rab18 binds to hepatitis C virus NS5A and promotes interaction between sites of viral replication and lipid droplets. PLoS Pathog 9, e1003513, doi:10.1371/journal.ppat.1003513 (2013).

39 Gerondopoulos, A., Bastos, R. N., Yoshimura, S., Anderson, R., Carpanini, S., Aligianis, I., Handley, M. T., Barr, F. A. Rab18 and a Rab18 GEF complex are required for normal ER structure. J Cell Biol 205, 707–720, doi:10.1083/jcb.201403026 (2014).

40 Zeng, Y. & Cullen, B. R. RNA interference in human cells is restricted to the cytoplasm. RNA 8, 855–860, doi:10.1017/s1355838202020071 (2002).

41 Stalder, L., Heusermann, W., Sokol, L., Trojer, D., Wirz, J., Hean, J., Fritzsche, A., Aeschimann, F., Pfanzagl, V., Basselet P. et al. The rough endoplasmatic reticulum is a central nucleation site of siRNA-mediated RNA silencing. EMBO J 32, 1115–1127, doi:10.1038/emboj.2013.52 (2013).

42 Pepin, G., Perron, M. P. & Provost, P. Regulation of human Dicer by the resident ER membrane protein CLIMP-63. Nucleic Acids Res 40, 11603–11617, doi:10.1093/nar/gks903 (2012).

43 Choudhary, R., Baturin, D., Fosmire, S., Freed, B. & Porter, C. C. Knockdown of HPRT for selection of genetically modified human hematopoietic progenitor cells. PLoS One 8, e59594, doi:10.1371/journal.pone.0059594 (2013).

