## Supplementary material for "*RAB18* is a key regulator of GalNAc conjugated siRNA induced silencing in Hep3B cells": All Supplementary Figures

| Cell Line | Aliases | Source | Tissue | Disease | ASGR1 [FPKQ] | ASGR2 [FPKQ] |
|---|---|---|---|---|---|---|
| HEPG2 | Hep G2 | CCLE | Liver | Hepatocellular Carcinoma | 297.5 | 203.7 |
| HEP3B217 | Hep 3B\|Hep 3B2.1-7\|Hep-3B\|Hep3B | CCLE | Liver | Hepatocellular Carcinoma | 135.9 | 208.9 |
| HUH7 | HuH-7 | CCLE | Liver | Hepatocellular Carcinoma | 65.5 | 175.3 |

Supplementary Table 1

| Molecule Type | siRNA ID | Gene Target | Conjugate format | Target mRNA Sequence |
|---|---|---|---|---|
| siRNA | 2035 | ASGR1 | GalNAc | 5’GUGGGAAGAAAGAUGAAGUUU3’ |
| siRNA | 1424 | ASGR1 | GalNAc | 5’GUGGGAAGAAAGAUGAAGUUU3’ |
| siRNA | 8172 | HPRT1 | GalNAc | 5’UCCUAUGACUGUAGAUUUUA3’ |
| siRNA | 17102 | HPRT1 | Cholesterol | 5’UCCUAUGACUGUAGAUUUUA3’ |
| siRNA | 6709 | HPRT1 | Anti-ASGR1 Antibody | 5’UCCUAUGACUGUAGAUUUUAU3’ |
| siRNA | 16084 | ASGR1 | GalNAc | 5’GGGAAGAAAGAUGAAGUCGC3’ |
| siRNA | 8714 | PPIB | GalNAc | 5’UUGGAAAGACUGUUCCAAAA3’ |

Supplementary Table 2

| gRNA Name | Sanger Clone ID | Target Gene | Species | DNA Target Sequence |
|---|---|---|---|---|
| SLC3A2-83 | HS5000011883 | SLC3A2 | Human | GCATGACTGGAGCCTCCATAGG |
| SLC3A2-84 | HS5000011884 | SLC3A2 | Human | CCAGCTTCCCTGACATCCCAGG |
| ASGR1-77 | HS5000003177 | ASGR1 | Human | TTCACGTGGAGCAGCAGGCTGG |
| ASGR1-78 | HS5000003178 | ASGR1 | Human | CGGAGCCTGAGCTGTCAGATGG |

Supplementary Table 3

| | Supplier: | Cat# | siRNA ID | Target | Sequence (sense) | Sequence (antisense) |
|---|---|---|---|---|---|---|
| **siRAB18-1** | Ambion | 4390824 | s22703 | Hs. RAB18 | GGAUGGAAAUAAGGCUAAAtt | UUUAGCCUUAUUUCCAUCCac |
| **siRAB18-2** | Ambion | 4390824 | s22704 | Hs. RAB18 | GGUUCACAGAUGAUACGUUtt | AACGUAUCAUCUGUGAACCtc |
| **siRAB18-3** | Ambion | 4390824 | s22705 | Hs. RAB18 | GGAAAAUCGUGAAGUCGAUtt | AUCGACUUCACGAUUUUCCtt |

Supplementary Table 4

a

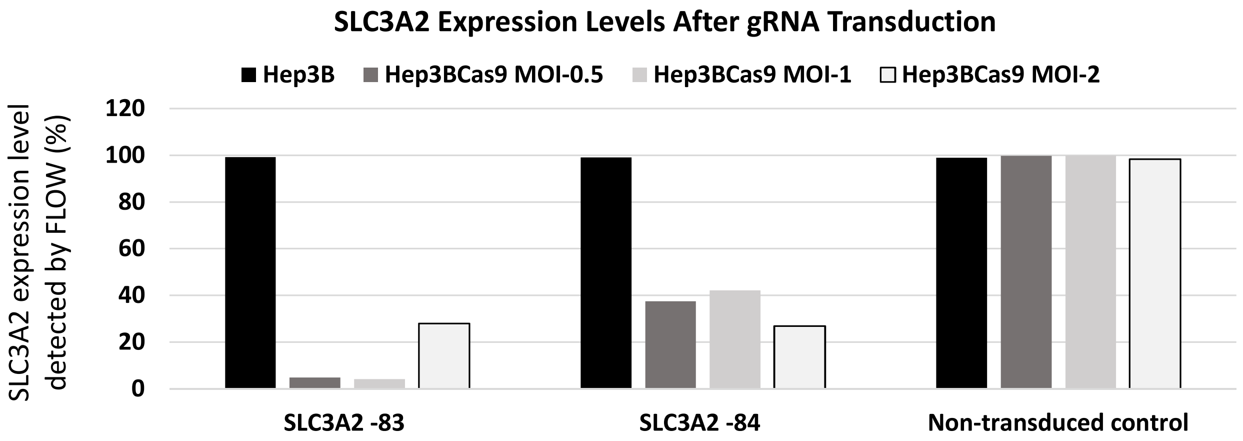


b

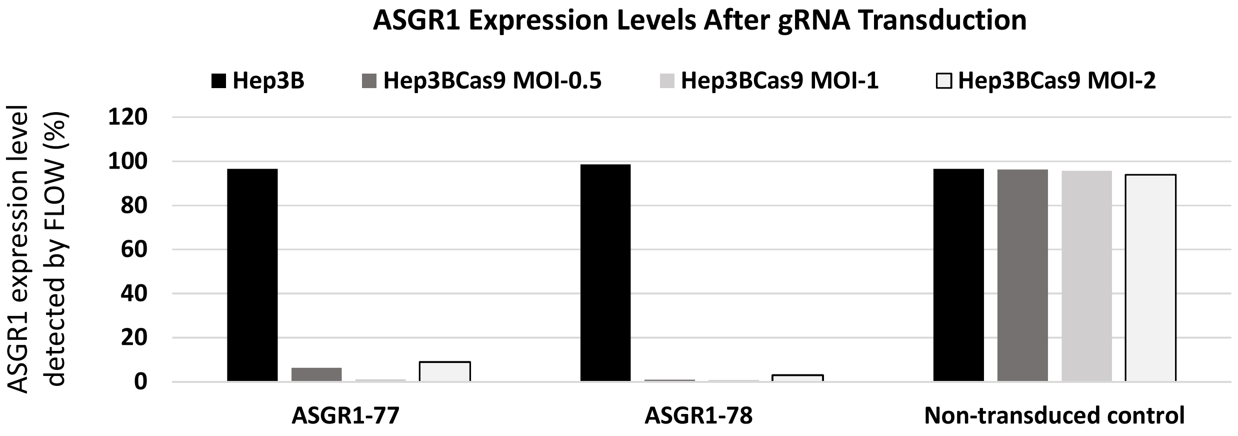


Supplementary Figure 1

### Validate HPRT1-si Resistance to 6TG in CRISPR Library Transduced Hep3BCas9 Cells

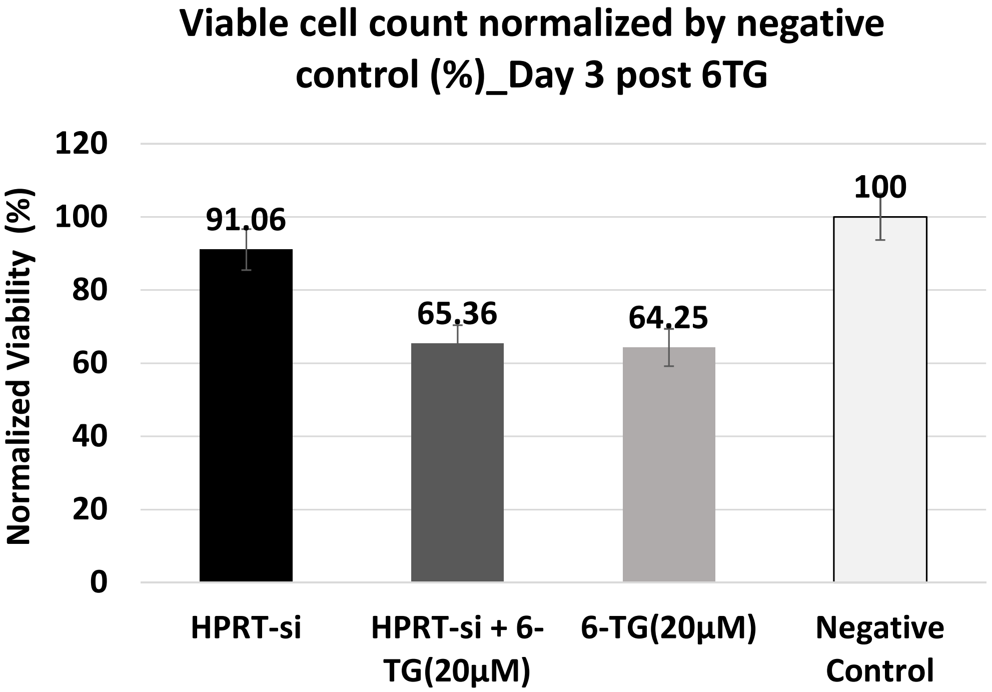


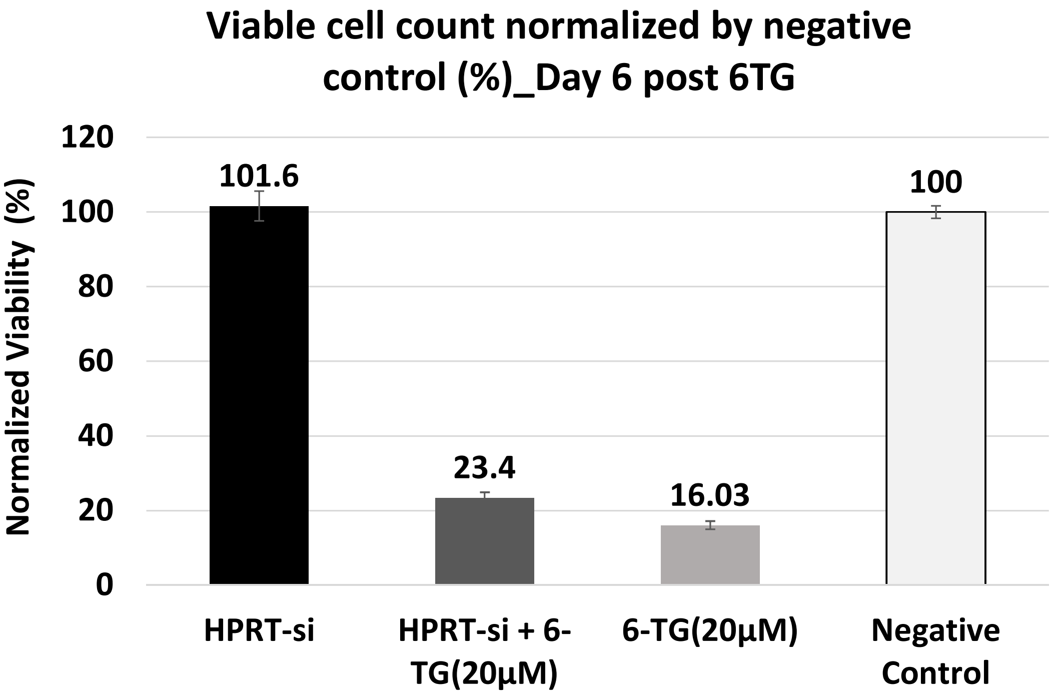


Supplementary Figure 2

a

| Sample Name | Description |
|---|---|
| plasmid | Plasmid library-sgRNA counts from sequencing plasmid library used to make lentiviral library for screen |
| BSL | Baseline sample |
| nosid3 | No siRNA treatment samples harvested on day 3 post siRNA treatment (no 6TG) |
| 150sid3 | 150nM siRNA treated samples harvested on day 3 post siRNA treatment (no 6TG) |
| 750sid3 | 750nM siRNA treated samples harvested on day 3 post siRNA treatment (no 6TG) |
| nosid9 | No siRNA treatment samples harvested on day 9 post siRNA treatment (no 6TG) |
| 150sid9 | 150nM siRNA treated samples harvested on day 9 post siRNA treatment (no 6TG) |
| 750sid9 | 750nM siRNA treated samples harvested on day 9 post siRNA treatment (no 6TG) |
| nosi6TGd9 | 6TG treated no siRNA treatment samples harvested on day 9 post siRNA treatment) |
| 150si6TGd9 | 6TG treated 150nM siRNA treated samples harvested on day 9 post siRNA treatment |
| 750si6TGd9 | 6TG treated 750nM siRNA treated samples harvested on day 9 post siRNA treatment |

b

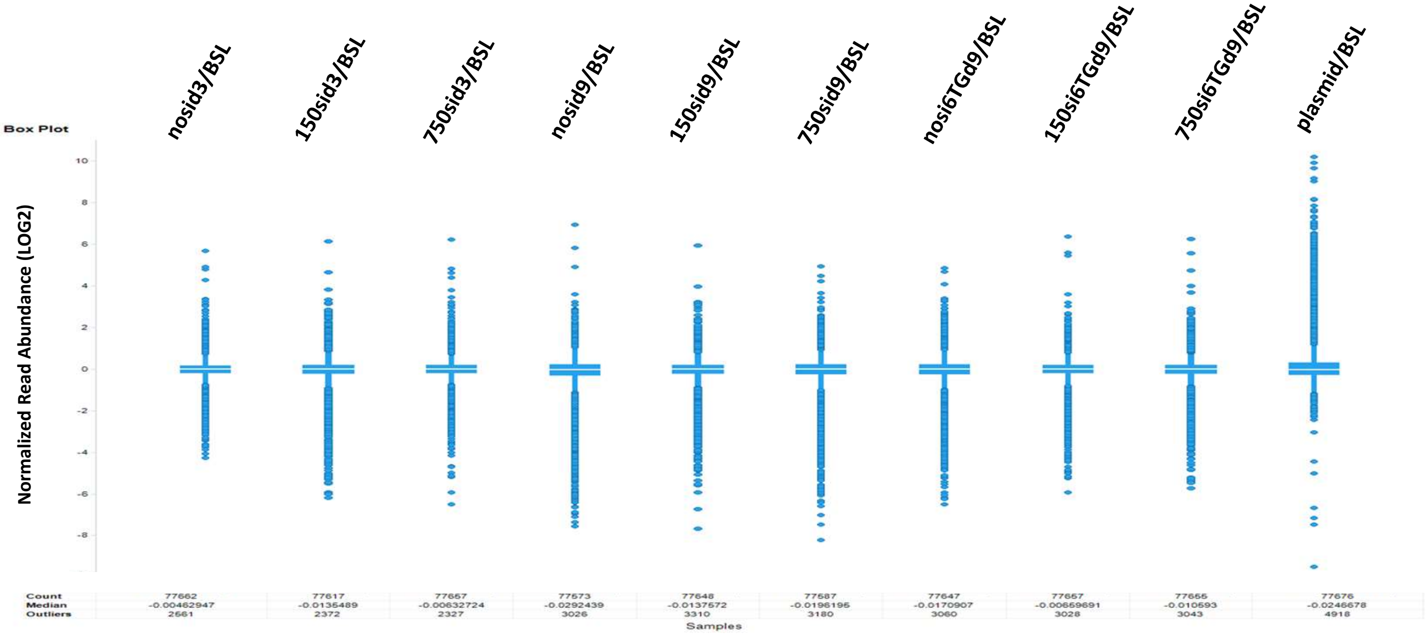


Supplementary Figure 3

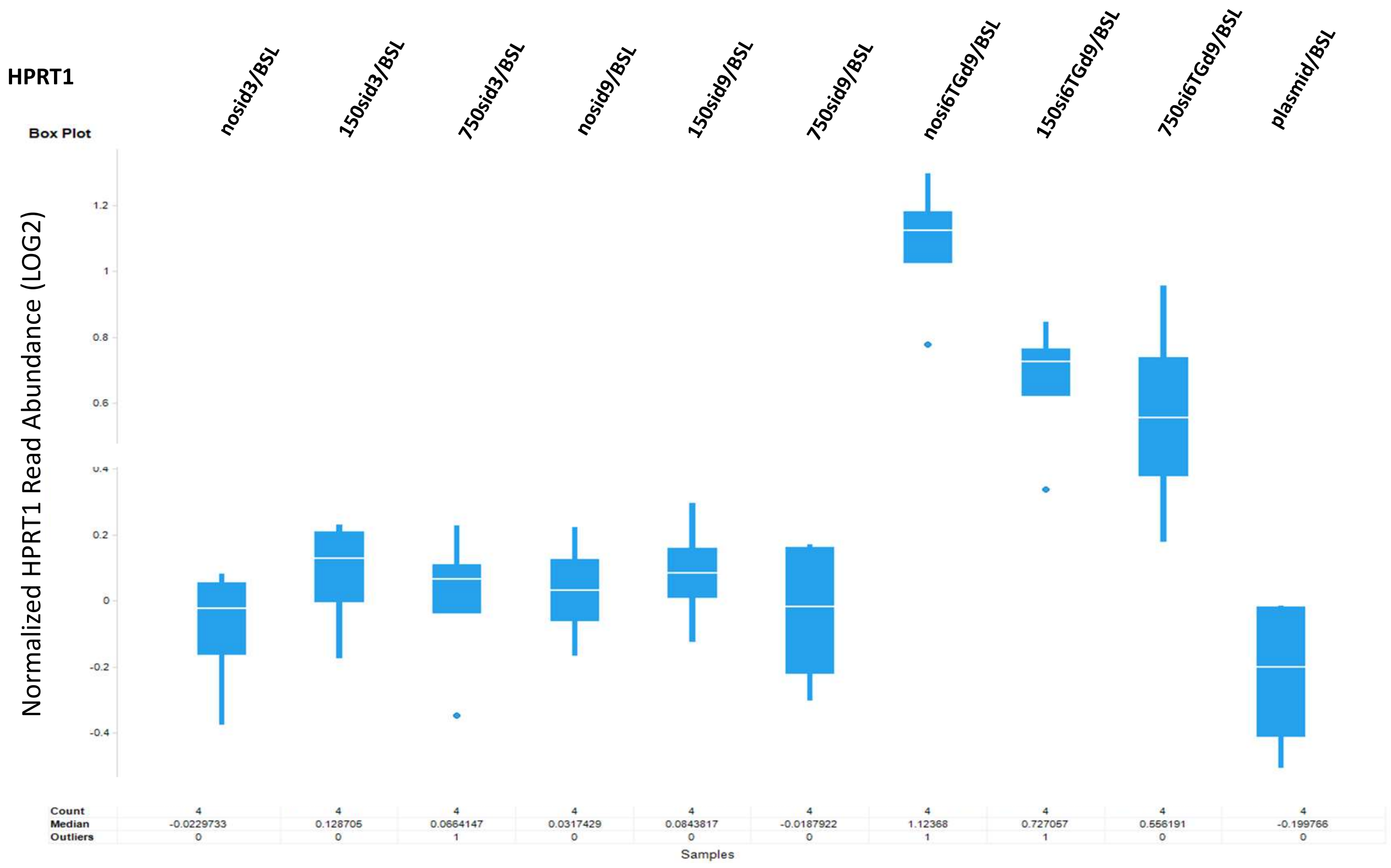


Supplementary Figure 4

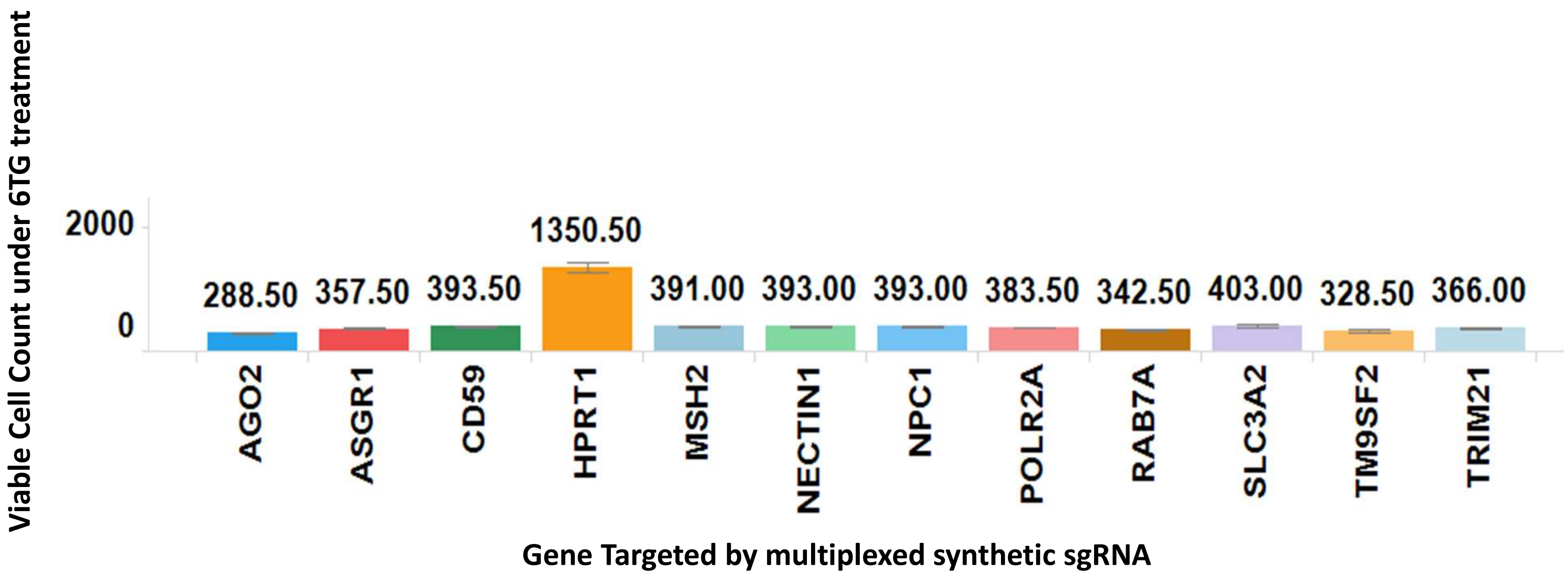


Supplementary Figure 5

a

SIGMA Sanger Vector:
U6-gRNA:PGK-puro-2A-tagBFP

| Sanger Clone ID | Gene Symbol | DNA Target Sequence |
|---|---|---|
| HS5000033611 | RAB18 | TAACTCCCAGCTATTATAGAGG |
| HS5000033612 | RAB18 | GCTATTATAGAGGTGCACAGGG |

b

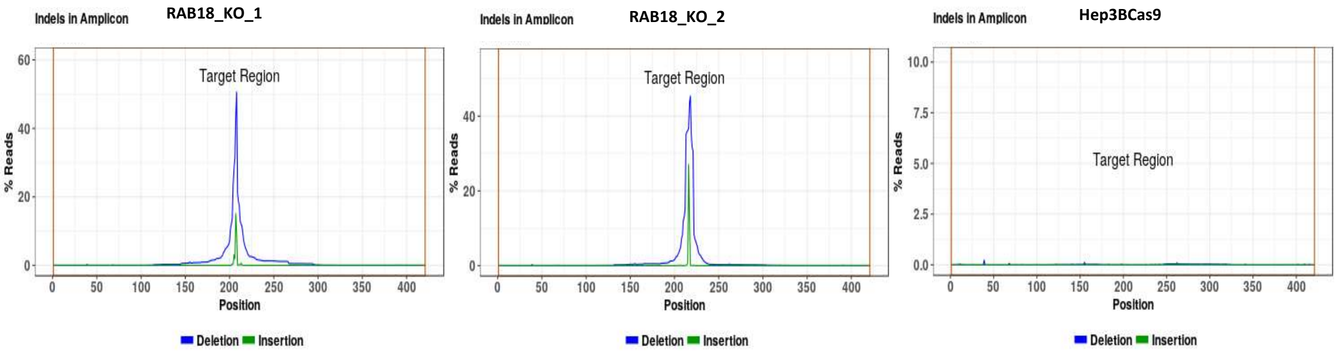


d

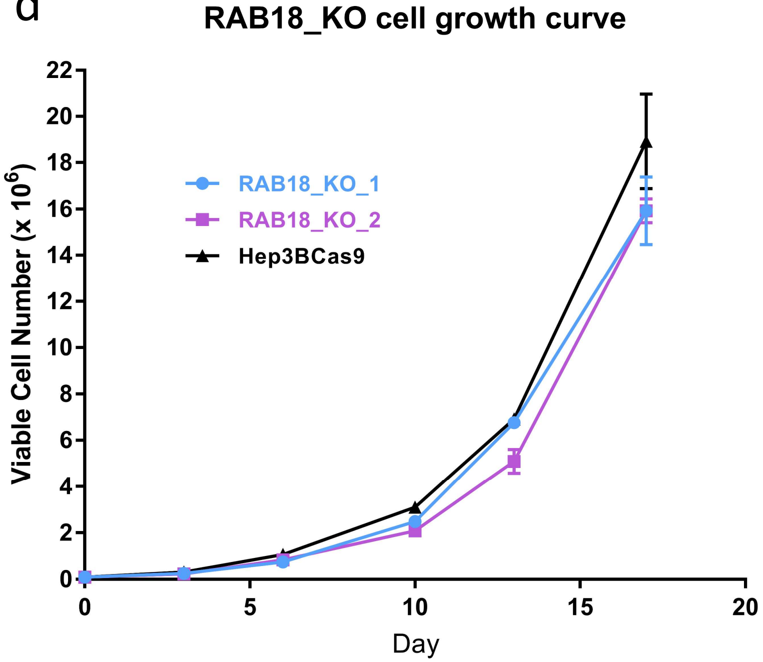


c

| Sample | Target Reads | Mutant Reads | Mutant Pct | Genotype | Frameshift Mutant Reads | Frameshift Mutant Pct |
|---|---|---|---|---|---|---|
| RAB18_KO_1 | 53595 | 52639 | 98.22 | Homozygous Mutant | 43884 | 81.88 |
| RAB18_KO_2 | 57155 | 57029 | 99.78 | Homozygous Mutant | 41116 | 71.94 |
| Hep3BCas9 | 53151 | 689 | 1.3 | Homozygous WT | 667 | 1.25 |

Supplementary Figure 6

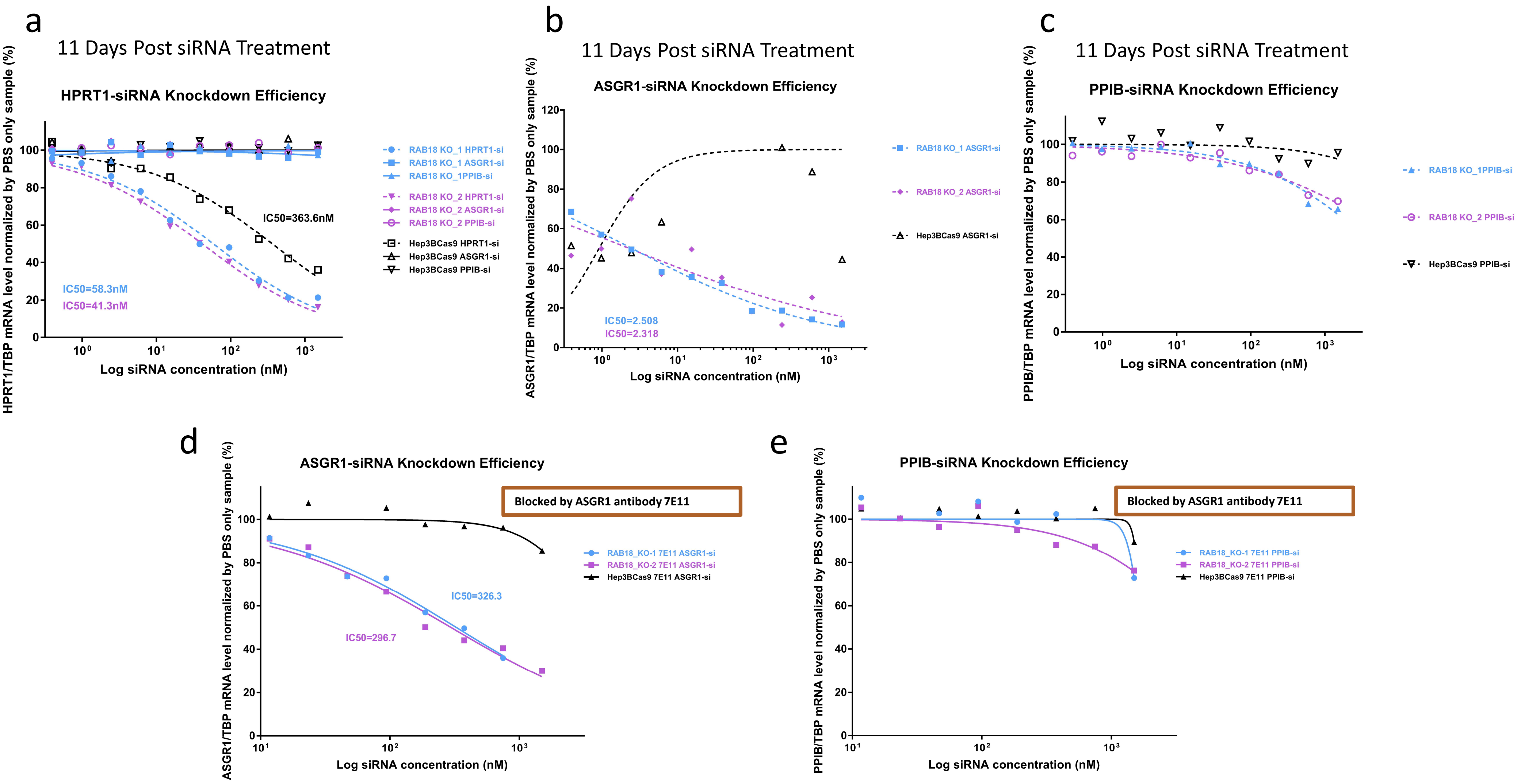


Supplementary Figure 7
